# Transcriptome and Comparative Genomics analyses reveal new functional insights on key determinants of pathogenesis and interbacterial competition in *Pectobacterium* and *Dickeya* spp

**DOI:** 10.1101/391854

**Authors:** Daniel Bellieny-Rabelo, Collins K. Tanui, Nikki Miguel, Stanford Kwenda, Divine Y. Shyntum, Lucy N. Moleleki

**Author notes:** Present address: Stanford Kwenda - Sydney Brenner Institute for Molecular Bioscience, University of the Witwatersrand, Johannesburg, Gauteng, South Africa. **Mesh Unique IDs:** D023281; D020869; D044043; D001435; D000069376; D016667.

## Abstract

<u>S</u>oft-<u>r</u>ot *<u>E</u>nterobacteriaceae* (SRE) typified by *Pectobacterium* and *Dickeya* genera are phytopathogenic bacteria inflicting soft-rot disease in crops worldwide. By combining genomic information from 100 SRE with whole-transcriptome datasets, we identified novel genomic and transcriptional associations amongst key pathogenicity themes in this group. Comparative genomics revealed solid linkage between the <u>t</u>ype <u>I</u> <u>s</u>ecretion <u>s</u>ystem (T1SS) and the carotovoricin bacteriophage (Ctv) conserved in 96.7% of *Pectobacterium* genomes. Moreover, their co-activation during infection might indicate a novel functional association involving T1SS/Ctv. Another bacteriophage-borne genomic region mostly confined in less than 10% of *Pectobacterium* organisms was found, presumably comprising a novel lineage-specific prophage in the genus. We also detected the transcriptional co-regulation of a previously predicted toxin/immunity pair (WHH and SMI1_KNR4 families) along with type VI secretion system (T6SS) and *hcp/vgrG* genes suggesting a role in disease development as T6SS-dependent effectors. Further, we showed that another predicted T6SS-dependent endonuclease (AHH-family) exhibited toxicity in ectopic expression assays indicating antibacterial activity. Additionally, we report the striking conservation of group-4-capsule (GFC) cluster in 100 SRE strains which consistently features adjacently conserved serotype-specific gene-arrays comprising a previously unknown organization in GFC clusters. Also, extensive sequence variations found in *gfcA* orthologs suggest a serotype-specific role in the GfcABCD machinery.

## 1. Introduction

Conflicts amongst microbes, and against their hosts, unfolds under intense selective pressure, which have selected an abundant inventory of colonization traits in unicellular organisms from all domains of life (Alouf, 2003, Aznar *et al*., 2015, Berg, 1975, Costerton *et al*., 1999, Gossani *et al*., 2014, Kubheka *et al*., 2013). In prokaryotic genomes specifically, the allocation of gene-inventories is recognized by the frequent conservation within functionally clustered units. Such pattern is determined by the strong selection imposed over the organization of prokaryotic genomes (Touchon & Rocha, 2016). The typical simultaneous occurrence of essential processes, such as transcription, translation and protein localization in the prokaryotic cytoplasm drives selective pressure on the genome organization (Esnault *et al*., 2007). In this scenario, the occurrence of clustered units constitutes a cohesive solution to facilitate co-transcription of functionally associated genes (Overbeek *et al*., 1999, Sabatti *et al*., 2002, Tamames *et al*., 1997, Yin *et al*., 2010). In fact, this feature has been broadly exploited through guilty-by-association methods in several studies, enabling the elucidation of novel gene functions (Anantharaman & Aravind, 2003, Overbeek *et al*., 1999, Zhang *et al*., 2011).

Bacteria from the *Enterobacteriaceae* family responsible for causing soft rot-blackleg/wilt disease are collectively designated as <u>S</u>oft-<u>R</u>ot *<u>E</u>nterobacteriaceae* (SRE) which are mainly represented by *Pectobacterium* and *Dickeya* genera (Pérombelon, 2002). These microbes have attracted attention for causing great impact on global food-security, as they infect a considerable range of plant hosts (Marquez-Villavicencio *et al*., 2011, Pérombelon, 2002). Hence, roughly one-hundred SRE complete-genome sequencing projects are publicly available, either completed or currently in progress (Alic *et al*., 2015, Li *et al*., 2015, Onkendi *et al*., 2016, Panda *et al*., 2015, Raoul des Essarts *et al*., 2015). Underpinned by such wealth of public data, recent transcriptome-based reports have depicted important facets of SRE biology. For example, the role of small RNAs in the adaptive response of *Pectobacterium atrosepticum* exposed to nutrient-deficient environments, revealing 68 regulated sRNAs in these conditions, and the discovery of nine novel sRNAs (Kwenda *et al*., 2016). Also in *P. atrosepticum*, the impact on the transcription of 26% of the genome upon deletion of *expI* revealed the critical role of quorum sensing regulation for disease development (Liu *et al*., 2008). Similarly, transcriptome analyses demonstrated the impact of 32 isolated stress conditions, mimicking those found during infection, on *Dickeya dadantii* strain 3937 regulatory patterns, providing a detailed landscape on environmental triggers for gene expression (Jiang *et al*., 2016). Furthermore, an investigation on the role played by the *D. dadantii* strain 3937 PecS global regulator during early colonization of leaf tissues uncovered more than 600 genes in its regulon (Pedron *et al*., 2017). Thus, as a general rule, transcriptome-based approaches are powerful means to explore both key strategies utilized by plant pathogens, and also plants’ critical physiological programs, which can potentially optimize plant genetic enhancement resulting in positive long-term impact on food-security (Bellieny-Rabelo *et al*., 2016, Gao *et al*., 2013, Tanui *et al*., 2017).

The deployment of several exoenzyme-families specialized on breaking plant cell wall is one of the most conspicuous strategies presented by SRE, hence one of the most exhaustively investigated (Toth & Birch, 2005). The activity of plant cell wall degrading enzymes (PCWDE) release byproducts which can be taken up as nutrients by the bacterial cell (Pérombelon, 2002). Some of these PCWDE include cellulases, pectate lyases (PLs) and pectin lyases (PNLs) amongst others (Allen *et al*., 1989, Pérombelon, 2002). Another important asset for disease development is the ability to biosynthesize high-molecular weight polysaccharides, which may either be secreted extracellularly (EPS - exopolysaccharide), or remain attached to the bacterial cell surface (e.g. lipopolysaccharide (LPS) and (CPS) capsular polysaccharides) (Whitfield, 2006). The expression of <u>h</u>orizontally <u>a</u>cquired <u>i</u>slands (HAIs) also comprise an ubiquitous strategy to induce pathogenesis and to succeed in interbacterial competition (Ochman *et al*., 2000). Genomically integrated bacteriophages (prophages) and toxins/antitoxins systems frequently exported by bacterial secretion systems (e.g. Type I, III, VI Secretion Systems) typify pathogenically important HAIs (Durand *et al*., 2014, Varani *et al*., 2013). Specifically, since the discovery of the bacteriophage-like <u>t</u>ype <u>VI</u> <u>s</u>ecretion <u>s</u>ystem (T6SS), it has been implicated as a crucial ecological asset of several Gram-negative bacteria either as a virulence or bacterial-competition agent (Pukatzki *et al*., 2006).

In this article regions of interest were gleaned by analyzing an original transcriptomic dataset obtained from *P. carotovorum* subsp. *brasiliense* strain PBR 1692 (henceforth referred to as *Pcb* 1692) during disease development. The aim was to survey the transcriptional activation of critical genomic regions required for virulence or interbacterial competition in *Pcb* 1692 and assess their conservation in 100 SRE genomes. We report herein the *in planta* co-activation of the <u>c</u>aro<u>t</u>o<u>v</u>oricin homolog (Ctv) prophage in *Pcb* 1692 along with a T1SS module immediately upstream. Comparative analysis added support to this evidence by unveiling an exquisite genomic conservation of the T1SS+Ctv block in 96.7% of all *Pectobacterium* strains analyzed. These results presumably point to a systemic association between these two themes in *Pectobacterium* genera, in which T1SS may export Ctv elements through the periplasm. Extensive gene neighborhood and protein domain-architecture analyses combined with large-scale sequencing also shed light, for the first time, to the strong topological and transcriptional association of a previously predicted toxin/immunity pair (WHH- and SMI1_KNR4-containing gene-products), to the T6SS machinery. Further evidence uncovered the up-regulation of ~71% (17 out of 24) of the genes in a ~25 kb region during the first 24 <u>h</u>ours <u>p</u>ost-<u>i</u>nfection (hpi) in *Pcb* 1692, which comprises a highly conserved capsule biosynthesis cluster in SRE. The analyses also demonstrated that the group <u>4</u> <u>c</u>apsule (GFC, or G4C) may be the only capsule production region conserved in *Pcb* 1692. Sequence analyses of gene-products encoded by *gfcA* locus in a large number of organisms unveiled high sequence variation, suggesting a putative role in the GFC machinery as a serotype-specific membrane protein.

## 2. Results and Discussion

### Transcriptome sequencing of *Pcb* 1692 during *in planta* infection

*Pcb* 1692 has been reported as one of the most aggressive *Pectobacterium* species known to date (Duarte *et al*., 2004, Durrant, 2016, Marquez-Villavicencio *et al*., 2011). Aiming to examine the transcriptional landscape of *Pcb* 1692 during infection in potato tubers, a whole-transcriptome dataset was generated including samples harvested 24 and 72 hpi along with *in vitro* control (see ‘Experimental Procedures’). The dataset comprises 14 to 20 million RNA-Seq paired-end reads for each stage, with ~97% of the reads in each sample uniquely mapped on the reference genome implying good overall quality (Table 1). Subsequent analyses identified 1743 protein-coding genes (43,5% of total annotated in *Pcb* 1692) under infection-induced regulation (log2fold-change > 1 or < −1; FDR < 0.01) in the wild-type strain in at least one time range (Table S2). Importantly, a recent study depicted *D. dadantii’*s transcriptome during infection on *A. thaliana*, compared at 6 and 24 hpi, being able to detect 13,5% of its protein-coding genes (575 out of 4244) under regulation (up or down) (Pedron *et al*., 2017). In the next steps, we take advantage of this recently published gene-expression experiment featuring a closely related wild-type SRE strain infecting a non-crop host, to comparatively examine up/down-regulation in our original gene-expression dataset obtained from *Pcb* 1692.

**Table 1.** Mapping summary of RNA-Seq reads on Pcb 1692 reference genome.

| Sample* | Unmapped Reads | Multiple Matches | Uniquely Mapped Reads | % Uniquely Mapped | Total Mapped reads | % Total Mapped | Raw Reads Inputs |
| --- | --- | --- | --- | --- | --- | --- | --- |
| *Cwt* | 751 568 | 416 981 | 13 835 241 | 97,07 | 14 252 222 | 95,0 | 15 003 790 |
| *Wt-24hpi* | 1 856 371 | 555 623 | 18 131 044 | 97,03 | 18 686 667 | 91,0 | 20 543 037 |
| *Wt-72hpi* | 2 852 242 | 333 569 | 11 124 857 | 97,09 | 11 458 426 | 80,1 | 14 310 667 |

\* Experimental samples are abbreviated as follows: Cwt - *in vitro* control wild-type; Wt-24hpi - wild-type 24 hours post infection; Wt-72hpi - wild-type 72 hours post-infection

SRE’s most well-recognized strategy to accomplish plant tissues invasion involves disruption of plant cell wall by deploying a variety of pectinase-, cellulase- and protease-related families (Pérombelon, 2002). PLs and PNL are pectinases that participate in this process by breaking α-1-4-linkages of homogalacturonan backbone in the pectin molecule via transelimination mechanism (Carpita & Gibeaut, 1993). The dataset presented here emphasizes the role of PL in *Pcb* 1692 infection by the significant activation of seven out of eight genes, occurring simultaneously during the first 24 hpi. This encompasses orthologs of well-documented genes (*pelABCINZ – PCBA_RS04055;04060;04065;10065;03200;04070*) and one yet unannotated gene (*PCBA_RS18630*) (Table S2). These findings are similar to those detected in *D. dadantii*, in which all nine PLs (*pelABCDEILNZ*) were activated while infecting *A. thaliana* (Pedron *et al*., 2017). The pectin lyase gene *pnl (PCBA_RS19200)*, which encodes a major PCWDE (Liu *et al*., 2008), is massively activated (> 5 log2fold-change) ranking amongst the top 1% most up-regulated genes 24 hpi in *Pcb* 1692 (Table S2). This finding suggests a specific high transcriptional demand of *pnl* in *Pcb* 1692 during early disease development. Conversely, in *D. dadantii* transcriptome, the *pnl* ortholog (*Dda3937_03551*) surprisingly shows no detectable regulation (Table S3) (Pedron *et al*., 2017). Together these results strengthen the notion of diversity in PCWDE pools transcriptionally activated by pectinolytic pathogens to surpass various obstacles imposed by hosts in distinct plant tissues (Toth *et al*., 2003). Whereas for some PCWDE families (e.g. PLs) the up-regulation is basal, regardless of host characteristics, other PCWDE families (e.g. PNL) conversely, are transcriptionally activated under specific host-imposed cues.

### Transcriptional profile and conservation of type VI secretion system and associated effectors in SRE

The T6SS is a bacteriophage-like structure that depends on the expression of 13 core genes (*tss* cluster) to assemble a complex that spans through the cell envelope (Pukatzki *et al*., 2006). The <u>h</u>aemolysin <u>c</u>orregulated <u>p</u>rotein (Hcp) forms a tube that is propelled across the membranes with a piercing structure on its tip, consisting primarily of two proteins (VgrG/PAAR), into the target cell (Basler, 2015, Shneider *et al*., 2013). Furthermore, it has also been demonstrated that this piercing structure is able to accommodate independent proteins which function as effectors (Durand *et al*., 2014). Our dataset underscores the importance of T6SS with the overwhelming activation of all 13 genes in tss cluster (> 3 log2fold-change in transcriptional variation during the first 24 hpi). Moreover, a total of four Hcp-secretion-islands (HSIs), one adjacent to tss cluster (HSI-1), and another three in different genomic contexts (HSI-2, -3 and -4), were also up-regulated in at least one time point. These HSI exhibit an overall activation of two PAAR-coding and four *vgrG* genes (Table S4). Comparatively, wild-type *D. dadantii* displays similar up-regulation of most tss and four *hcp* genes out of six conserved in the genome upon infection on *A. thaliana* (Table S4). However, none of the VgrG- or PAAR-encoding genes (five and three respectively) are transcriptionally regulated in *D. dadantii* in the same experiment (Pedron *et al*., 2017). Notably, the role of VgrG-PAAR as a delivery mechanism for Rhs effectors into target cells is established in *D. dadantii* facing *in vitro* competition (Koskiniemi *et al*., 2013). Thus, although structural components of T6SS sheath and tube are similarly regulated when infecting distant hosts in *Pcb* 1692 and *D. dadantii*, genes encoding the piercing tip components may be activated under specific cues (Pedron *et al*., 2017) (Table S3).

Interestingly, the second major HSI in *Pcb* 1692 genome (HSI-2) spans ~6.6 kb harboring eight genes under positive regulation in at least one time point during infection (Fig. 1A). Of these, six (excluding only *hcp* and *vgrG*) are uncharacterized/unannotated (Table S4).

**Figure 1.**
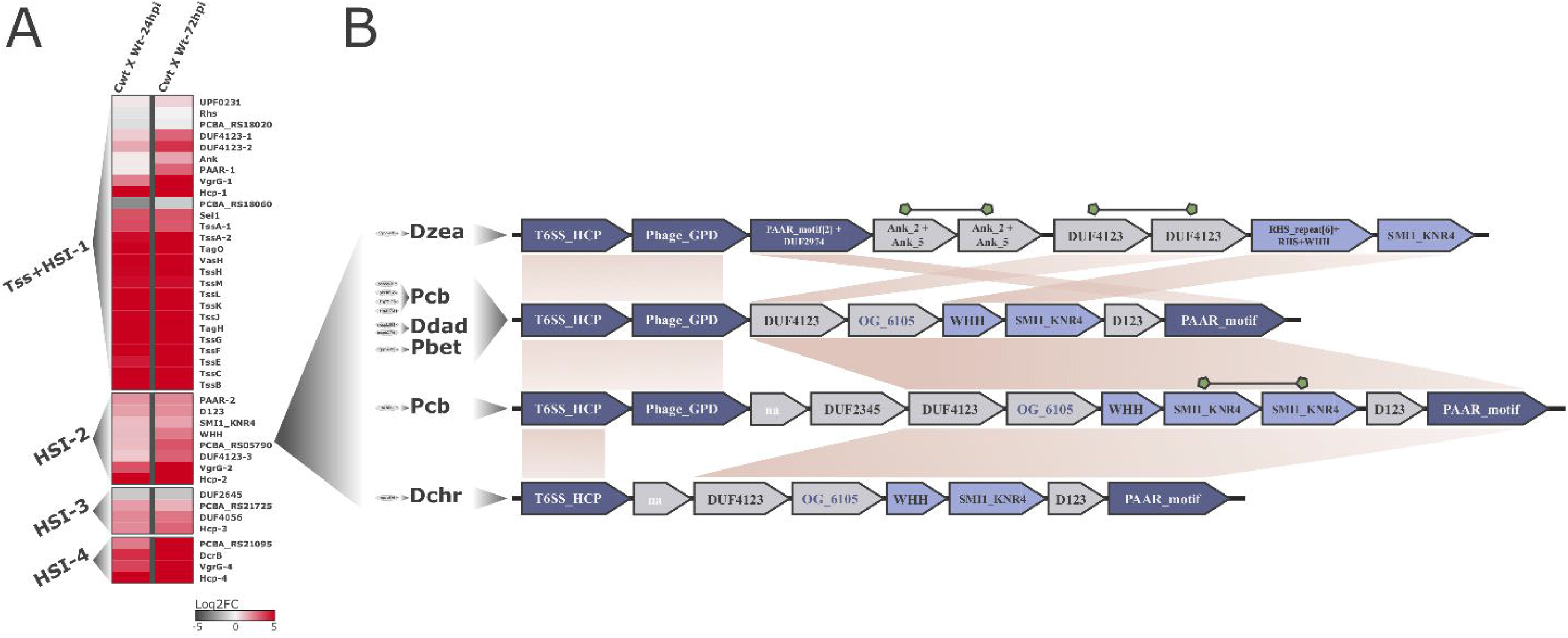
Transcriptional variation *in planta* of T6SS-related genes in *Pcb* 1692 and linkage between the endonuclease WHH and HSI in SRE genomes: **(A)** Each row on the heatmap represents a gene labeled after its (i) respective encoded-protein name, (ii) functional domain or (iii) locus-tag respectively given the annotation availability. The two columns of the heatmaps represent pairwise comparisons between samples, thus each cell shows the variation in transcription found in each comparison illustrated by a color-scale ranging from -5 (dark-grey) to 5 (red) representing the log2fold-change variation in expression. Column labels are abbreviated as follows: Cwt = Control wild-type; Wt-24hpi/Wt-72hpi = wild-type 24/72 hours post-infection in potato tubers. Four segments in the heatmap separate T6SS-related clusters - type <u>VI</u> secretion (Tss) and Hcp secretion island (HSI) - located in different genomic regions, namely: Tss+HSI-1, HSI-2, -3 and -4. **(B)** Gene neighborhood of WHH-SMI1_KNR4 duplets in the 10 SRE strains conserved within HSIs. Each gene arrangement is associated with its respective strain(s) represented on the left side by 4-letters abbreviations as follows: *Dzea (Dickeya zeae), Pcb (Pectobacterium carotovorum* subsp. *brasiliense), Ddad (Dickeya dadantii), Pbet (Pectobacterium betavasculorum), Dchr (Dickeya chrysanthemi)*. For each species, the respective number of strains conserving that particular gene arrangement are represented in grey ellipses on their left. The WHH-SMI1_KNR4 duplets (Pfam: PF14414; PF09346 respectively) are highlighted in light-blue. Hcp (T6SS_HCP; Pfam: PF05638), VgrG (Phage_GPD; Pfam: PF05954) and PAAR (PAAR_motif; Pfam: PF05488) encoding genes are highlighted in dark-blue. Syntenically conserved blocks across the different gene arrangements are linked by light-brown areas. Duplicated genes in a single arrangement are linked by green shapes. Successfully clustered products in orthologous-groups for which no conserved domains were detected are highlighted in dark-blue font.

Hence, aiming to garner deeper functional insight on this region, we combined analysis of domain architectures conserved in protein sequences with contextual genomic information from 100 SRE organisms. Firstly, conserved domains analysis confirmed VgrG (PCBA_RS05800; PFAM: PF05954) and PAAR gene-products (PCBA_RS05770; PFAM: PF05488) in this region, implying that the cluster is able to source both Hcp-tube, and the piercing tip (VgrG/PAAR) assembly (Fig. 1B). Further, two uncharacterized genes (*PCBA_RS05780; PCBA_RS05785*) flanked by the genes encoding VgrG and PAAR caught our attention for containing respectively SMI1_KNR4 and WHH (PFAM: PF14414 and PF09346) domain architectures in their products. Interestingly, the association between the superfamilies SUKH and HNH/ENDOVII, which encompass SMI1_KNR4 and WHH families respectively, has been recognized as a recurrent theme in bacterial genomes (Zhang *et al*., 2011). This duplet (WHH-SMI1_KNR4) was described as an immunity/toxin pair associated with contact-dependent growth inhibition (CDI) systems (e.g. T5SS, T6SS), although evidence actually linking these genes to the secretion systems remains scarce (Aoki *et al*., 2010, Hayes *et al*., 2010, Ruhe *et al*., 2013, Zhang *et al*., 2011). The WHH family was originally identified as a restriction endonuclease highly derived from the HNH domain (PFAM: PF01844; CL0263) (Shub *et al*., 1994). Our analyses revealed that members of the WHH family spread across 18 out of 100 SRE strains. Of these, tight association to SMI1_KNR4 family-members immediately downstream in 13 genomes was found, which corroborates the original report for this duplet (Fig. 1B and Table S4) (Zhang *et al*., 2011). Out of the 13 genomes conserving this association (WHH-SMI1_KNR4) in SRE, one was not suitable for contextual genomic inspection in this specific region due to incompleteness of genome assembly (namely *P. betavasculorum* strain NCPPB 2793). By assessing the remaining suitable structures conserving WHH-SMI1_KNR4 duplet, we found 83% (10 out of 12) linkage with upstream HCP, and PAAR-encoding genes mostly downstream (Fig. 1B). A duplication in the immunity gene encoding the SMI1_KNR4 was also found in one strain, which corroborates the original report of this family (Zhang *et al*., 2011) (Fig. 1B). These results imply a strong association of this duplet with HSIs in SRE genomes, such as: *Pcbs* (five strains including *Pcb* 1692), *D. dadantii* (two strains), *D. zeae* (one strain), *D. chrysanthemi* (one strain) and *P. betavasculorum* (one strain) (Fig. 1B and Table S4). These observations taken together with the coordinated up-regulation at the transcriptional level of WHH-SMI1_KNR4 encoding genes along with all other HSI-2 neighboring elements in *Pcb* 1692 suggest the T6SS-dependent secretion of these genes (Fig. 1). In this context, we provide the first report of coordinated transcriptional regulation of a SUKH-1/HNH-ENDOVII system along with the surrounding HSI, in which the WHH-containing gene-product may work as a T6SS-effector recruited during infection.

### Analysis of antibacterial activity of type VI secretion system related toxins

Aiming to investigate the potential of HSI-borne genes from *Pcb* 1692 to have antibacterial activity, four relevant genes were cloned into an expression vector and ectopically expressed in *E. coli*, as described by Koskiniemi *et al*. (2013). To this end, the previously, described WHH-containing gene (PCBA_RS05785) was selected alongside three other putative toxins (PCBA_RS05775, PCBA_RS18045 and PCBA_RS22965) for this analysis. PCBA_RS05775 belongs to the cell division cycle protein 123 family (D123; Pfam: PF07065), which seems not to be accompanied by adjacent immunity encoding gene (Table S4). The PCBA_RS18045 gene is located in HSI-1, within the *Pcb* 1692 tss gene cluster and possesses a PAAR domain followed by an alpha/beta hydrolase fold with a GxSxG catalytic lipase motif (Pfam: PF12697), characteristic of phospholipases (Alcoforado Diniz *et al*., 2015). This phospholipase family member is followed by a putative immunity gene downstream that carries an Ank domain (Pfam: PF00023). The PCBA_RS22965 gene has a conserved HNH/EndoVII fold derivative known as AHH (Pfam: PF14412). Similarly to WHH-family members, the AHH proteins were predicted to be functionally associated with CDI systems in the original report of the family (Zhang *et al*., 2011). In most SRE genomes harboring members of the AHH family, no detectable domain in the downstream immunity protein can be observed (Table S4). In *Pcb* 1692, similarly to six other SRE genomes, a contig break occurs up to five genes prior to this AHH-member, hindering direct gene neighborhood based conclusions. Nonetheless, in other strains from *Pcb, Pcc* and *Patr* species conserving 70-95% of overall genomic synteny with *Pcb* 1692 (Fig. S1) the AHH genes are solidly neighbored by *hcp, vgrG* and a DcrB domain (Pfam: PF08786) encoding gene, suggesting a role as a T6SS-associated effector (Table S4). Such arrangement precisely matches the previously assessed HSI-4 region in *Pcb* 1692 (Fig. 1A). In addition to the original report of AHH family, these results suggest that the AHH member probably play a role in *Pcb* 1692 infection as a T6SS-associated nuclease

The results show that no cell death occurred due to expression of three of the four effectors between 60 and 240 minutes (Fig. 2A). However, expression of the AHH effector, caused a reduction in *E. coli* growth, indicating that it has a toxic effect on *E. coli*. This growth inhibition could be counteracted by co-expression of the toxin and immunity gene (AHH+i) in *E. coli*, suggesting that the previously observed reduced growth was due to the expression of the AHH nuclease (Fig. 2B). Thus far, no experimental data exists for the AHH and WHH nucleases, as well as the D123 protein as T6SS effectors. This is the first experimental work featuring the antibacterial activity of a AHH-containing protein.

**Figure 2.**
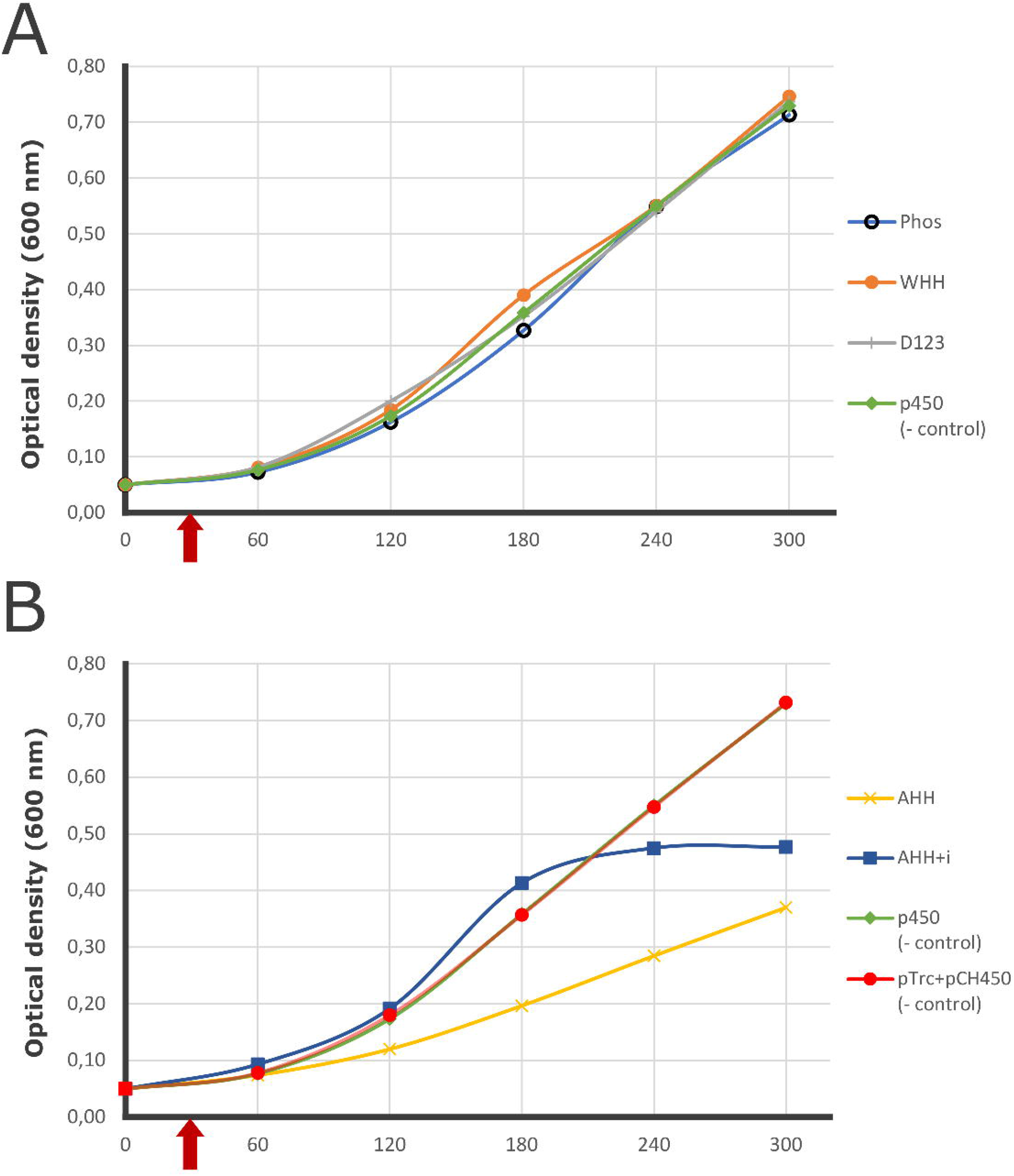
Growth of *E. coli* DH5-alpha expressing effectors from pCH450 induced with 0.2% L-arabinose after 30 min growth, or both effector and immunity protein from pCH450 and pTrc100, respectively. **(A)** Expression of AHH nuclease decreases the growth rate of *E. coli*, but not as pronounced as the positive control RhsA from *D. dadantii* (Koskiniemi *et al*., 2013). Expression of both AHH and its immunity protein Ai (AHH +i) negate toxic effects of the effector alone, however after extended periods displays growth inhibition due to protein overexpression. **(B)** Expression of phospholipase, WHH nuclease, and D123 effectors grows comparable to the empty vector control (p450), indicating that individual expression of these effectors does not contribute towards bacterial killing in the condition analyzed.

Several T6SS effectors targeting eukaryotes, bacteria, or both have been identified thus far (Jiang *et al*., 2016, Pukatzki *et al*., 2007, Russell *et al*., 2011). Here, we have demonstrated that one of the four effectors identified have a killing effect on *E. coli* cells. It is possible that, for the other effectors tested, the *in vitro* conditions used were not conducive for observable killing. It may also need to be considered that our hypothesis that the T6SS of *Pcb* 1692 is mainly antibacterial may not be entirely correct. Bernal *et al*. (2018) have noted that thus far, no plant-targeting effectors have been identified. As the D123 protein has no reported function and lacks a downstream immunity protein, this protein could be a candidate for a plant-targeted effector. Since some effectors have inter-kingdom activity, it may be necessary to re-evaluate whether the effectors we have identified as putative antibacterial effectors may have key functions within the eukaryotic host. This is the first experimental work featuring the antibacterial activity of a AHH-containing protein. Moreover, it must also be taken into consideration that T6SS could have functions other than contact-dependent antibacterial competition which may favour conservation of effectors involved in other biological programs that will enforce successful host colonization.

### Genomic conservation and transcriptional regulation of prophages during *in planta* infection

The impact of HAIs originating from prophages in bacterial pathogenic behavior is pervasive for both plant and animal pathogens (Addy *et al*., 2012, Reeve & Shaw, 1979, Vaca-Pacheco *et al*., 1999). Importantly, selective advantage conferred by prophage-borne genes to plant-pathogens has been associated with the activity of bacterial secretion systems, as reported in *Pseudomonas syringae* and *Ralstonia solanacearum* (Genin & Denny, 2012, Guidot *et al*., 2007, Varani *et al*., 2013). In this context, by using a combination of relevant genome-wide approaches (see ‘Experimental Procedures’), we detected two prominent prophage regions in *Pcb* 1692 exhibiting significant activation at the transcriptional level during infection (Fig. 3A). The first region encompasses 20 genes in *Pcb* 1692 genome remarkably up-regulated in the first 24 hpi (Fig. 3A). Moreover 90% of these genes are syntenic with genomic segments in at least 54/60 *Pectobacterium* strains (Fig. 3B and Table S5). Interestingly, a genomic segment in *P. carotovorum* subsp. *carotovorum* (*Pcc*) syntenic to the one found in *Pcb* 1692 was originally described as of prophage origin, regarded as <u>c</u>aro<u>t</u>o<u>v</u>oricin (Ctv) consisting of 19 genes (Yamada *et al*., 2006). The report of this bacteriocin has been recently corroborated by *in silico* analyses in the same species, even extending the putative range of the cluster to 22 genes in *Pcc* (termed PecaPC1-p2) (Table S5) (Varani *et al*., 2013). In this context, we also detected in *P. atrosepticum* (*Patr*) a segment of 12.7 kb containing 11 gene-products highly similar to Ctv sequences (ECA_RS18415-18485). Intriguingly, this segment is located within a previously characterized bacteriophage spanning ~36 kb, designated ECA41 (Evans *et al*., 2010). In addition, ECA41 contribution to virulence towards potato hosts was reported, although the mechanism of putative effectors’ action remains unknown (Evans *et al*., 2010). Curiously, those three reports describing Ctv, PecaPC1-p2 and ECA41 occurred without knowledge convergence based on the detectable homology linking the regions in *Patr* and *Pcc*, which our analyses now reconcile under the same Ctv-homologous root. This preliminary observation may provide a robust background for future studies on carotovoricin role in pathogenesis with the combined interpretation of the above-mentioned reports.

**Figure 3.**
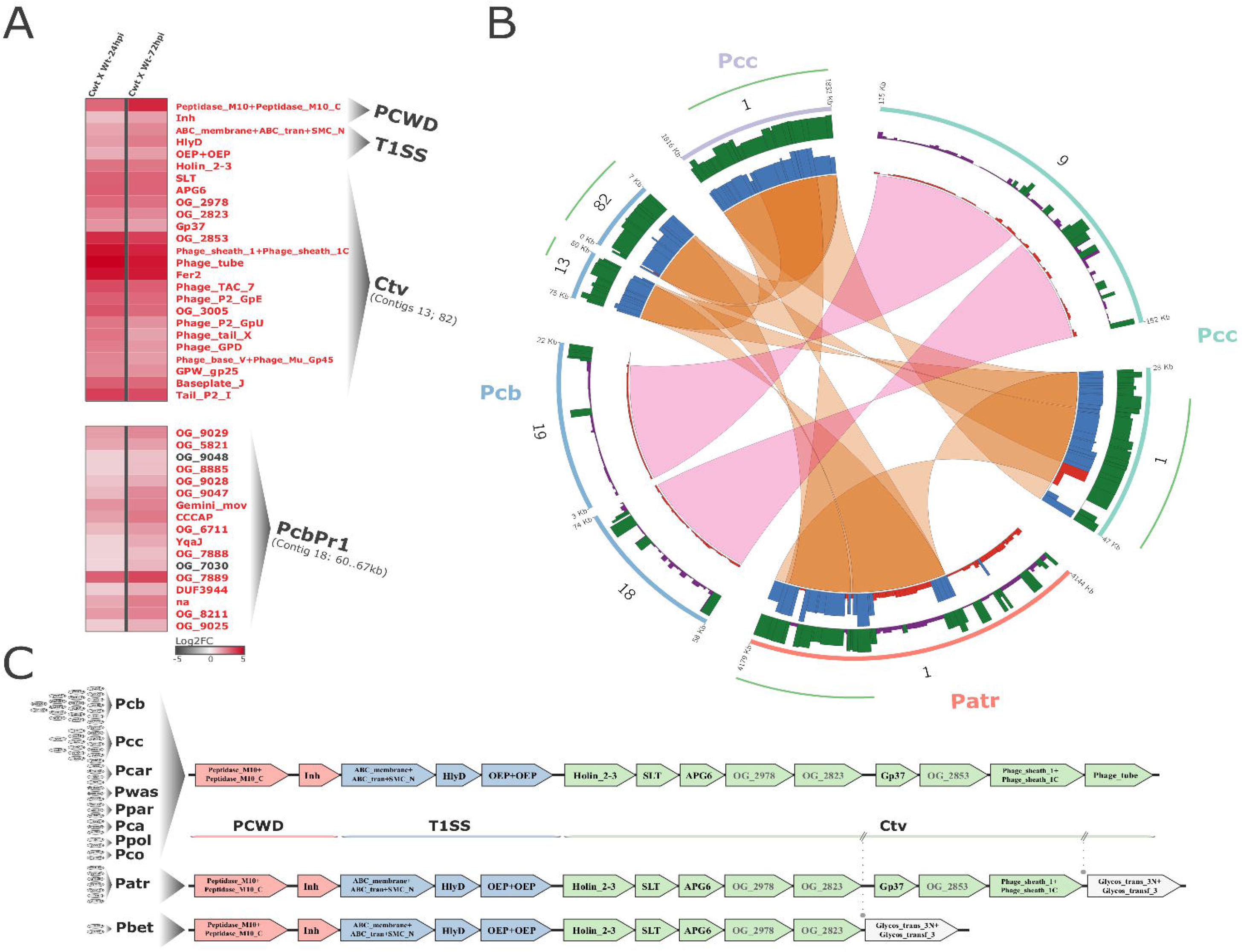
Transcriptional variation in *Pcb* 1692 and genomic range of Carotovoricin (Ctv) and PcbPr1 insertions in four *Pectobacterium* strains. **(A)** The heatmaps show transcriptional variation profiles of T1SS+Ctv block and serralysin/inhibitor genes (top) and from a 7 kb segment of PcbPr1 (bottom). The heatmap is represented as in Figure 1. For Ctv and PcbPr1, contig coordinates are represented in parenthesis immediately below labels, which links to the item B of this panel. Genes are preferentially represented by the domain architecture, or by the respective orthologous group (OG_#). The “na” label is used if none of the previous is applicable. Genes in red font are significantly up-regulated, whereas grey indicates no support for regulation. **(B)** Ctv and PcbPr1 drastically contrasting degree of conservation is represented in four selected *Pectobacterium* strains. Genomic regions from *Pcc* strain PC1 (light-purple) and BCS2 (light-green), *Patr* strain SCRI1043 (light-orange), and *Pcb* 1692 (light-blue) are represented in circular ideograms. The next two consecutive inner radiuses respectively represent the frequencies of Blastp (protein sequence similarity – purple→green range) and MCScanX (syntenic conservation – red→blue range) pairwise hits of each *Pcb* 1692 sequence compared to 99 SRE strains. The color-range key used in the bars represents the frequency of positive hits found for each *Pcb* 1692 sequence: <= 30 (purple/red) or > 30 (green/blue). The links binding genomic regions represent the insertion range of prophages for which conserved synteny could be detected (Ctv = orange; PcbPr1 = pink). The green outer-radius adjacent to the ideograms represent the range of Blastp similarity with the original Ctv described by Yamada *et al*. (2006). (Table S5) **(C)** Gene neighborhood screening of T1SS+Ctv block: Functional groups are separated in: plant cell wall degradation (PCWD - light-red), T1SS (blue) and Ctv (light-green) or unrelated to these classes (grey). Each genomic organization found in *Pectobacterium* genomes is sided by the species conserving a respective arrangement abbreviated as in Table S9. Each species is sided by grey ellipses representing the respective number of strains harboring the particular arrangement.

Furthermore, we observed that Ctv is neighbored by a complete set of T1SS genes in *Pcb* (*PCBA_RS08610-08620*) immediately upstream (Fig. 3C and Table S5). In addition, conserved domains analyses revealed two genes in Ctv cluster containing SLT (PFAM: PF01464) and Gp37 (PFAM: PF09646) architectures, both reportedly conserved in bacterial virulence factors (Mushegian *et al*., 1996, Summer *et al*., 2004). Therefore, the presence of predicted effector functionalities in these genes along with a known secretion system prompted us to assess the degree of conservation of T1SS+Ctv structure. The results support a remarkable conservation of this genomic architecture, displaying a prominent block of at least 13 genes (Fig. 3C). This includes (a) two genes comprising a peptidase/inhibitor pair, (b) three T1SS components and (c) eight Ctv genes, under solid conservation in 96.7% of *Pectobacterium* genomes (59 out of 61). Importantly, this high conservation trend seems to occur regardless of the overall synteny between these genomes (Fig. S1 and Table S5). The T1SS is recognized for its relative simplicity, composed of three core proteins which form a tunnel-like structure in the inner membrane enabling molecule transfer from cytosol to the extracellular space fueled by ATP hydrolysis (Holland *et al*., 2005). Although typically regarded as signal-independent system, there is a known group of products exported in a T1SS-dependent manner that contains N-terminal signals. This group encompasses bacteriocins and microcins (Duquesne *et al*., 2007a, Duquesne *et al*., 2007b, Kanonenberg *et al*., 2013). Since Ctv constitutes a bacteriocin system, an additional layer of support to the T1SS/Ctv functional association could be added by searching for signal peptides within Ctv sequences. Given the generally poor conservation described in T1SS signal peptides, sensitive search setups were carried out (Emanuelsson *et al*., 2007, Frank & Sippl, 2008). The predictions from two different methods corroborate the possible T1SS/Ctv association by the combined detection of N-terminal signal peptides in five proteins encoded in this region (Supporting Information 1-2). In addition, some genes lacking known conserved domains, for which the protein sequences were clustered respectively in the <u>o</u>rthologous-<u>g</u>roups (see ‘Experimental Procedures’) OG_2978, OG_2823, and OG_2853, are conserved in 96-100% of *Pectobacterium* genomes (Fig. 3C and Table S5). These highly conserved genes in *Pectobacterium* curiously encode small products (up to 109 aa) and could undertake a role in bacterial competition as small antimicrobial peptides. Here we unraveled the strikingly dominant theme encompassing T1SS and Ctv clusters in *Pectobacterium* genomes tied by different evidence sources. This linkage suggests that either (a) Ctv may comprise an addictive selfish element colonizing these genomes, or (b) similarly to other reported T1SS/bacteriocins associations, this system might have been recruited in *Pectobacterium* lineage to export Ctv-borne products either towards or through the cell membranes.

The second prophage, which apparently has not been described thus far, conserves low similarity with other SRE genomes being mostly confined into *Pcb* strains (henceforth referred to as PcbPr1) and one *Pcc* strain (Fig. 3B). Blastp searches of PcbPr1 sequences resulted in five proteins out of 64 (7.8%) returning matches against at least 90 SRE species (out of 99) (Fig. 3B). In contrast, overall 82.3% of *Pcb* 1692 protein coding sequences (3376 out of 4099) successfully match against at least 90 SRE (Table S6). The low conservation of PcbPr1 is also observable in terms of genomic organization, as 93.7% of the whole PcbPr1 structure (60 out of 64 genes) conserves synteny with less than 10% of all *Pectobacterium* genomes analyzed (Fig. 3B and Table S6). Despite the low degree of conservation, PcbPr1 harbors an internal segment spanning 7 kb, in which 88.2% of the genes (15 out of 17) are cohesively up-regulated over the first 72 hpi (Fig. 3A and Table S6). Importantly, 12 out of 17 genes in this region encode proteins for which known conserved domains could not be detected. In addition, most of these gene products were clustered in small orthologous groups with sequences from other *Pcb* and *Pcc* implying lineage-specific origin (Fig. 3A). This observation supports the previous synteny analysis, reinforcing the possibility of PcbPr1 comprising a lineage-specific insertion in *Pcb* and *Pcc*. Notably, genomic conservation of functional lineage-specific prophages is observable in both animal and plant pathogens as typified respectively in studies using *E. coli* and *R. solanacearum* (Gabriel *et al*., 2006, Guidot *et al*., 2009, Perna *et al*., 2001). Interestingly, it has been reported that two HAIs in *R. solanacearum*, namely *Rasop1* and *Rasop2*, showed no similarity with other phages previously sequenced in the species (Varani *et al*., 2013). In order to assess gene expression patterns of *Rasop1/2* during infection, we gleaned data from an existing whole-transcriptome experiment (Jacobs *et al*., 2012). Up-regulation of 58.6% and 62.5% of *Rasop1* (27 out of 46) and *Rasop2* (20 out of 32) genes was detected upon infection on tomatoes (Table S3) (Jacobs *et al*., 2012). The evidence presented underscores lineage-specific insertions conservation as an important strategy for some organisms, probably providing competitive advantage against close species not bearing the same traits. Therefore, PcbPr1 conservation and up-regulation during infection might have implications on *Pcb* 1692 success when facing *in planta* competition against other *Pectobacterium spp*. As one of the few relatively large lineage-specific genomic regions compared to other *Pectobacterium spp*, we speculate that PcbPr1 could be a key asset for the reported high level of virulence observed in Pcb.

### Characterization of dedicated polysaccharide-biosynthetic clusters

The composition of serotype-specific polysaccharide gene clusters gives rise to biochemical diversity in bacterial cell-surfaces (Schmid *et al*., 2015). The fundamental role of cell surface in virulence has been well established in both animal and plant pathogens (Barras *et al*., 1994, Kao & Sequeira, 1991, Rocchetta *et al*., 1999, Whitfield, 2006). In SRE, however, aside from the establishment of *E. coli rffG* homolog in *P. atrosepticum* as a probable player in enterobacterial common antigen biosynthesis (Toth *et al*., 1999), the overall knowledge on this subject remains limited. In this context, our dataset enabled detection of a 25 kb region in *Pcb* 1692 functionally associated with polysaccharide biosynthesis exhibiting significant co-activation of 17 out of 24 genes in the first 24 hpi (Fig. 4A). Through orthology-based annotation of *Pcb* 1692 genes compared with model organisms (see ‘Experimental Procedures’), and subsequent integration with STRING correlational database (von Mering *et al*., 2005), most of the gene-products were annotated into the LPS biosynthetic pathway (Fig. 4B and Table S7). Out of the remaining eight unannotated entries, seven were successfully characterized by conserved domains inspection through HMM-profiles (Eddy, 2009, Finn *et al*., 2010). Thus, additionally revealing well-known domains in polysaccharide-production such as: an integral membrane polysaccharide-specific transporter (PST; PFAM: PF14667), an acetyltransferase (AT; PFAM: PF13302), a cupin-like protein (PFAM: PF05523), and four glycosyltransferases (GT; PFAM Clans: CL0110; CL0111) (Fig. 4A and Table S7). The remaining unannotated gene (*PCBA_RS09180*) is related to *gfcA/yjbE* (Ferrieres *et al*., 2007, Peleg *et al*., 2005) described in *E. coli* and will be discussed in detail in the next section. As a general rule all these eight entries unsurprisingly display high sequence variation mostly comprising weakly conserved blocks amongst SRE (Fig. 4A). These lineage-specific genes presumably comprise a serotype-specific block in SRE polysaccharide biosynthesis.

**Figure 4.**
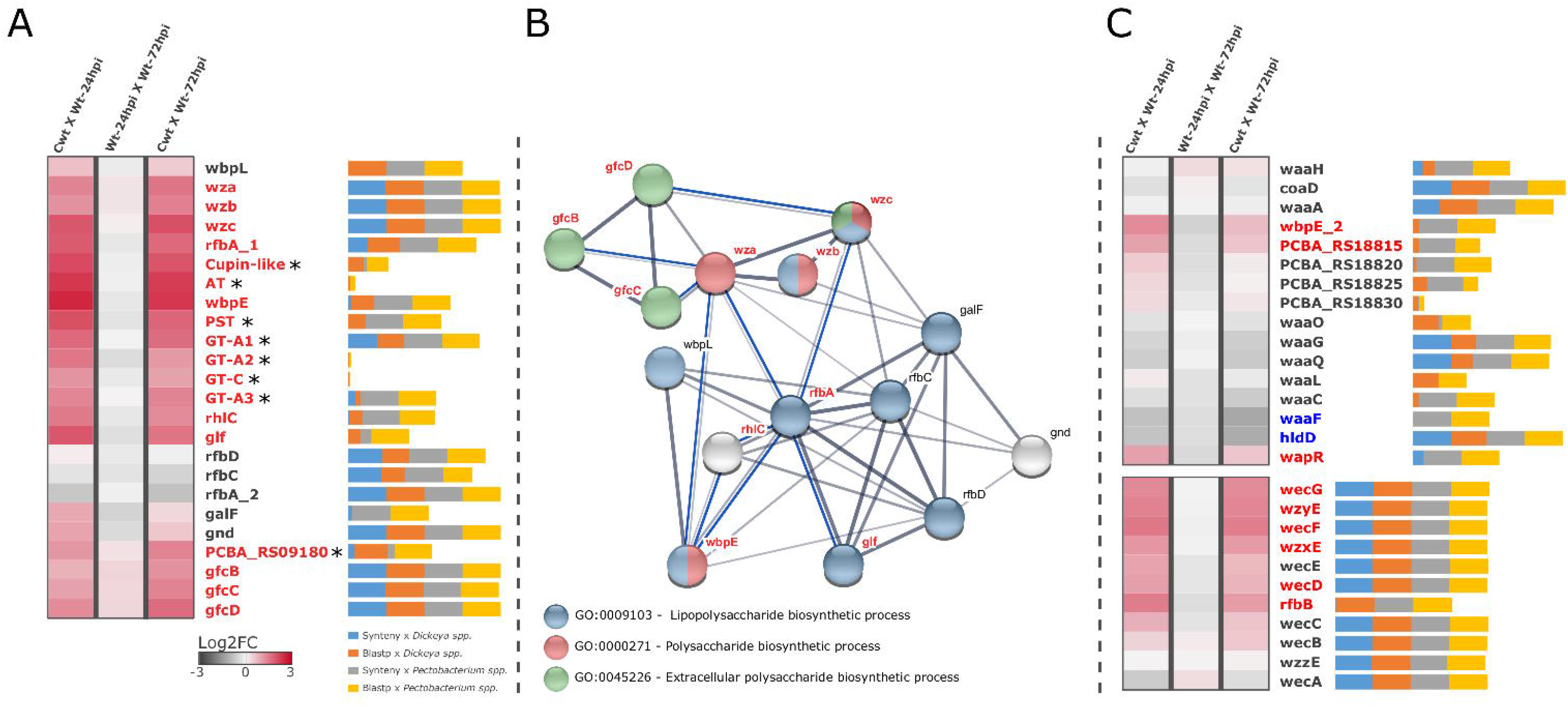
*In planta* transcriptional profile, genomic conservation and annotation of dedicated polysaccharide clusters in *Pcb* 1692: **(A)** Heatmap of GFC operon as represented in Figure 1. Genes highlighted with red font are significantly up-regulated in at least one interval. Asterisks next to the genes represent those for which no orthology relationship could be assessed, which by extension are not represented in the network. Next to each gene-name represented in the rows, horizontal bars (percentage-stacked) display their respective conservation compared with SRE genomes of *Dickeya* (39 strains) and *Pectobacterium* (60 strains) genera. These bars include overall frequencies of Blastp (sequence similarity) and MCScanX (syntenic conservation) searches of *Pcb* 1692 gene products against 99 SRE complete datasets. The bars are divided in segments: Syntenic with *Dickeya spp*. (blue), Blastp positive against *Dickeya spp*. (red), Syntenic with *Pectobacterium spp*. (grey), Blastp positive against *Pectobacterium spp*. (yellow). **(B)** Correlational network denotes the annotated gene association between *Pcb* 1692 successfully predicted orthologs in *E. coli* and *P. aeruginosa* according to STRING database (von Mering *et al*., 2005) also showing respective GO terms annotations. Thickness of network links are proportional to the combined scores obtained from STRING algorithm, which computes combined probabilities gleaned from different evidence sources listed in the tabular output found in (Table S7) (von Mering *et al*., 2005). Blue lines represent new co-xregulation evidence found in our transcriptome dataset between genes for which other categories of associations are known. Those entries not annotated in the three major GO terms shown in the network are highlighted in grey. **(C)** Transcriptional variation of *waa* (top) and *wec* (bottom) gene clusters represented as in (A). Additionally, down-regulated genes are highlighted in blue.

Next, by integrating our annotations with STRING networks, we observed new co-expression (coordinately up-regulated in our dataset) associations between *Pcb* 1692 orthologs during the course of infection (Fig. 4B and Table S7). The co-expression linking several genes encoding GTs and modification enzymes (e.g. *rhlC, rfbA, glf*, and *wbpE*) in our dataset were undetected in STRING, possibly indicating an uncommon genomic architecture in *Pcb* 1692 to be detailed below (Fig. 4B). Some other associations are surprisingly novel, such as those amongst *gfcBCD* and *wza/wzc* genes, all reportedly participants of capsule biosynthesis (Drummelsmith & Whitfield, 1999, Peleg *et al*., 2005) (Fig. 4B). In this regard, functional studies have attributed to Wza/Wzc the assembly of a transmembrane complex spanning the entire periplasm required for capsule assembly (Collins *et al*., 2007). As for *gfcABCD*, although the exact functions remain unknown, these genes are regarded as the typifying theme of group 4 capsule production clusters (Peleg *et al*., 2005, Rendueles *et al*., 2017, Whitfield, 2006). Therefore, the presence of *wza-b-c* and *gfc* orthologs indicates that *Pcb* 1692 possesses, and significantly activates during infection, the genetic apparatus to produce GFC (Fig. 4B and Table S7).

It is noteworthy that a recently proposed model for programmatic large-scale identification of GFC operons based on *E. coli* did not include internal serotype-specific genes (Rendueles *et al*., 2017). Indeed, a typical GFC operon in *E. coli* carries the *gfc* genes immediately upstream of *wza-b-c* paralogs (i.e. *gfcE, etp*, and *etk*) with no adjacent serotype-specific genes. However, our report supports the presence of lineage-specific genes flanked by *wza-b-c* and *gfcBCD* in *Pcb* 1692 for which transcriptional up-regulation is required upon infection, potentially representing a serotype-specific block (Fig. 4A). Moreover, the presence of lowly conserved gene organizations upstream of *gfc* genes seems to be a general feature in SRE genomes as depicted by extensive gene neighborhood analysis (Fig. S2 and Table S7). These blocks in SRE may vary from 13 to 27 genes harboring diverse family-compositions consistently flanked by *wza-b-c* upstream and *gfcBCD* downstream. Although blocks ranging from 16 to 20 serotype-specific genes seem to be preferred across SRE genomes, there is no direct association between any particular species and specific block sizes (Fig. S2 and Table S7). This ‘unusual’ organization of the GFC lineage-specific region in SRE compared to the one in *E. coli* might explain why some co-expression associations (e.g. *gnd*and *rhlC*) in our dataset were not present in STRING database as mentioned above (Fig. 4B). The possible functional implications of SRE and specially *Pcb* 1692 harboring this lineage-specific gene-array flanked by *wza-b-c* and *gfcABCD* will be discussed in detail in another section.

Importantly, a recent study predicted that 40% of the bacterial lineages analyzed conserve more than one capsule system (Rendueles *et al*., 2017). This observation prompted us to systematically survey *Pcb* 1692 genome for other possible capsule regions occurrences. Since LPS, EPS and capsules share many gene families in their functional units, which greatly hinders fully automatic predictions, the regions in *Pcb* 1692 genome were manually inspected following an initial automatic screening (see ‘Experimental Procedures’). The analysis allowed additional detection of two conspicuous polysaccharide clusters in *Pcb* 1692, although neither of these regions conserve characteristic domains that typify capsule-related regions (Table S7). Thus, these findings indicate that GFC may be the only capsule group produced in *Pcb* 1692. Nonetheless, these two additional regions carry respectively eight and seven *waa* and *wec* orthologs, which are commonly associated with LPS biosynthesis, and could be important assets during infection and therefore worth investigating (Lehrer *et al*., 2007, Regue *et al*., 2001). By analyzing their transcriptional patterns, an increased demand for GFC and *wec* genes transcription in the first 24 hpi was observed. Contrarily, in the *waa* region, only three out of 16 genes were up-regulated (Fig. 4C). Further, significant down-regulation of two genes (*waaF* and *hldD*), and three borderline predictions of down-regulation (*waaG, waaQ*, and waaC; Table S7) indicated a general low demand for transcription of the *waa* region during infection. Curiously, between 24-72 hpi a slight negative transcriptional modulation occurs in most of the genes in the three regions, however it does not significantly impact the overall trend for up-regulated genes (Fig. 4C). Unlike GFC or *waa* gene clusters, the *wec* region is remarkably conserved in nearly all SRE, encompassing six (out of 11) genes displaying infection-induced up-regulation in *Pcb* 1692 (Fig. 4C). These observations elucidate the unequal transcriptional demand for three distinct polysaccharide clusters during infection of potato by *Pcb* 1692. While *wec* and specially *gfc* gene clusters seem to be consistently recruited at the transcriptional level, the transcription profile of the *waa* cluster is mostly flat, suggesting different functional demands during infection for these regions.

### Analysis of *gfcA-related* sequences and genomic contexts

The inability to assess either orthology or domain conservation for PCBA_RS09180, in addition to the current lack of information regarding the *gfcA* function prompted an in-depth comparative investigation into this locus. Based on results presented above, 88% of the SRE analyzed genomes carry ‘orphan’ (not-clustered by OrthoMCL) genes upstream of *gfcBCD* (Table S7). In the remaining 22% strains, we detected gene products from small clusters populated with sequences from three (OG_5444: *D. solani, D. dadantii* and *D. chrysanthemi)*, two (OG_10199: *D. solani*, and *D. chrysanthemi)*, and one (OG_7594: *D. dianthicola*) species (Table S7). This clustering pattern in groups populated by a small number of species imply lineage-specific expansions. Furthermore, overall only 10% of SRE strains carry the YjbE domain (PFAM: PF11106) described in *E. coli* (Table S7). These include the two *P. betavasculorum* analyzed and six out of seven *P. atrosepticum* strains in the analysis. Hence, the presence of some PCBA_RS09180-related sequences in small clusters represented by no more than three species (e.g. OG_5444 and OG_7594) due to high level of sequence variation, constitutes a typical serotype-specific pattern.

Next, in order to test these results in a broader scope we expanded the gene neighborhood analysis beyond of the SRE group. Sequence-based searches were performed in order to retrieve a set of PCBA_RS09180-related proteins in bacteria. Unsurprisingly, Blastp (Altschul *et al*., 1990) search using PCBA_RS09180 sequence against NCBI (non-redundant protein database) was ineffective. In a preliminary level, this is consistent with the observations found in SRE, that showed high sequence variation in PCBA_RS09180-related sequences. We then utilized a more sensitive approach (Remmert *et al*., 2011) to obtain distantly related PCBA_RS09180 proteins. Within 50 positive matches, 27 were supported by publicly available genome-wide data, hence suitable for gene neighborhood screening (Table S7). From these, three encode YjbE-containing products (PFAM: PF11106) corroborating the relationship between *PCBA_RS09180* and *yjbE/gfcA*. On the other hand, the remaining 24 gene-products display no detectable domain (similarly to PCBA_RS09180). The results revealed that even in evolutionary distant organisms, 91% (22 out of 24) of the PCBA_RS09180 distantly related encoding genes are confined upstream of *gfc/yjb*-homologous operons (Table S7). Thus, although this locus has been termed *yjbE/gfcA* and the respective domain described in *E. coli* assigned as YjbE, this locus is in fact highly variable and the YjbE domain is weakly represented. An additional confirmation was obtained by consulting a large repository for domain architectures annotation (Geer *et al*., 2002). The presence of the YjbE domain in bacteria is at least 20-fold less frequent compared to YjbF, YjbH and Caps_synth_GfcC (Table S7). Together these results uncover the highly variable nature of the *gfcA* locus in a broad range of prokaryotes, which could be a serotype-specific player in the capsule biosynthesis machinery.

Next, the *gfcA* encoded products from 100 SRE strains were aligned in order to assess possible conserved residues in their sequences. The results revealed a conspicuously conserved N-terminal segment harboring a MKKTLxxLxxxxAxxxxxxA motif (Fig. S3).

Further, we conducted predictions for secondary structure of PCBA_RS09180 sequence in comparison to GfcA *and* YjbE using two different methods (Hofmann, 1993, Moller *et al*., 2001). Together, the analyses cohesively show two possible transmembrane sections in both PCBA_RS09180 and GfcA with apparent cytoplasmic orientation of N-terminal region (Fig. S4). Conversely, secondary-structure predictions for YjbE were inconclusive. Also, the prediction of N-terminal signal-peptide (Emanuelsson *et al*., 2007, Frank & Sippl, 2008) in PCBA_RS09180 and GfcA also strengthens the possibility of shared functionality (Supporting Information 3). In addition, it indicates closer functional relationship between PCBA_RS09180 and GfcA, compared to YjbE. Strikingly, further extensive signal peptide predictions made by two different methods in 100 *gfcA* sequences from SRE also returned 96% overlapping positive results in N-terminal regions (Supporting Information 4). This result matches the conserved region found in the previous alignment, reinforcing their functional conservation (Fig. S3). Moreover, secondary structure predictions in the same set of sequences supported that 80% of *gfcA* orthologs may conserve between one and three transmembrane regions, which is in accordance with the results from GfcA (Supporting Information 5). Together these results suggest that PCBA_ RS09180, and presumably SRE orthologs, could be directed to the inner-membrane (similarly to GfcA) to function as membrane proteins. This corroborates the current hypothesis on GfcABCD performing auxiliary role in polysaccharide translocation across the membranes (Sathiyamoorthy *et al*., 2011).

### Analysis of GFC elements and Wza/Wzc system organization in *Pcb* 1692

The currently accepted model for polysaccharide biosynthesis via Wza/Wzc transmembrane conduit involves association of several functional modules such as: (i) a series of peripheral GTs and sugar-modification enzymes, (ii) a Wzx-like (PST family) transmembrane flipping protein, (iii) an initiation sugar transferase, and (iv) a Wzy-like polymerase (Whitfield, 2006). The Wza/Wzc system is often associated with two highly similar systems that produce respectively group 1 and 4 capsules (Whitfield, 2006). Curiously, the gene arrangement in SRE’s GFC region resembles the one from group 1 capsule described in other species carrying a serotype-specific block downstream of *wza-b-c* (Fig. S2) (Rahn & Whitfield, 2003). Nonetheless, the genome-wide absence of the group 1 capsule-characteristic *wzi* gene suggests it could not be produced in the vast majority of SRE species, except for *P. betavasculorum*, and *P. carotovorum* subsp. *actinidiae* (Table S7). In this context, the results in *Pcb* 1692 were examined in the light of comparative analyses with SRE and then superimposed to the canonical Wza/Wzc model in order to propose a possible organization for the GFC machinery in *Pcb*. A total of eight GTs or modification enzymes, predicted as soluble or weakly attached to the inner membrane, encoded in this region exhibit cohesive up-regulation in the first 24 hpi in *Pcb* 1692 (Fig. 5). These enzymes exhibit 0-2 transmembrane (TM) sections, which is in accordance with the Wza/Wzc functional model (Fig. 5; Supporting Information 7 and Fig. S5). Next, the conserved PST within the GFC region presents 12 TM helices, which is described as the consensus number of TM sections for Wzx-like flippases (Islam & Lam, 2013). Also, the functional domain detected in this sequence is classified in the same Pfam clan (CL0222) which also encompasses the domain found in the canonical Wzx (Pfam: PF01943). Further, in *Pcb* 1692, the Glycos_transf_4 (PFAM: PF00953) domain found in the WbpL ortholog (PCBA_RS09080) is also conserved in WbaP/WecA sequences, which are the reported initiation transferase linked to Wzc in other bacteria (Table S7) (Valvano, 2003, Wang *et al*., 1996). This WbpL ortholog also features 11 TM helices solidly predicted, which matches the exact number found in WecA from *E. coli* (Fig. 5; Supporting Information 7 and Fig. S5). As for the Wzy-like activity, however, the direct association to this *Pcb* 1692 region is unclear. The WzyE activity in *E. coli* is described as similar to those performed by members of GT-B subfamily (Islam & Lam, 2014, Zhao *et al*., 2014). However, GT-B is the only unrepresented GT subfamily within the described GFC region in *Pcb* 1692. Interestingly, a GT-C found within the *Pcb* 1692 GFC region conserves 10 TM helices, matching the number found in *Pcb’s* direct ortholog of WzyE (PCBA_RS14975). The presence of periplasmic loops could also be predicted in this GT-C member, consistent to previously described Wzy proteins (Fig. 5 and Supporting Information 7) (Daniels *et al*., 1998, Kim *et al*., 2010).

**Figure 5.**
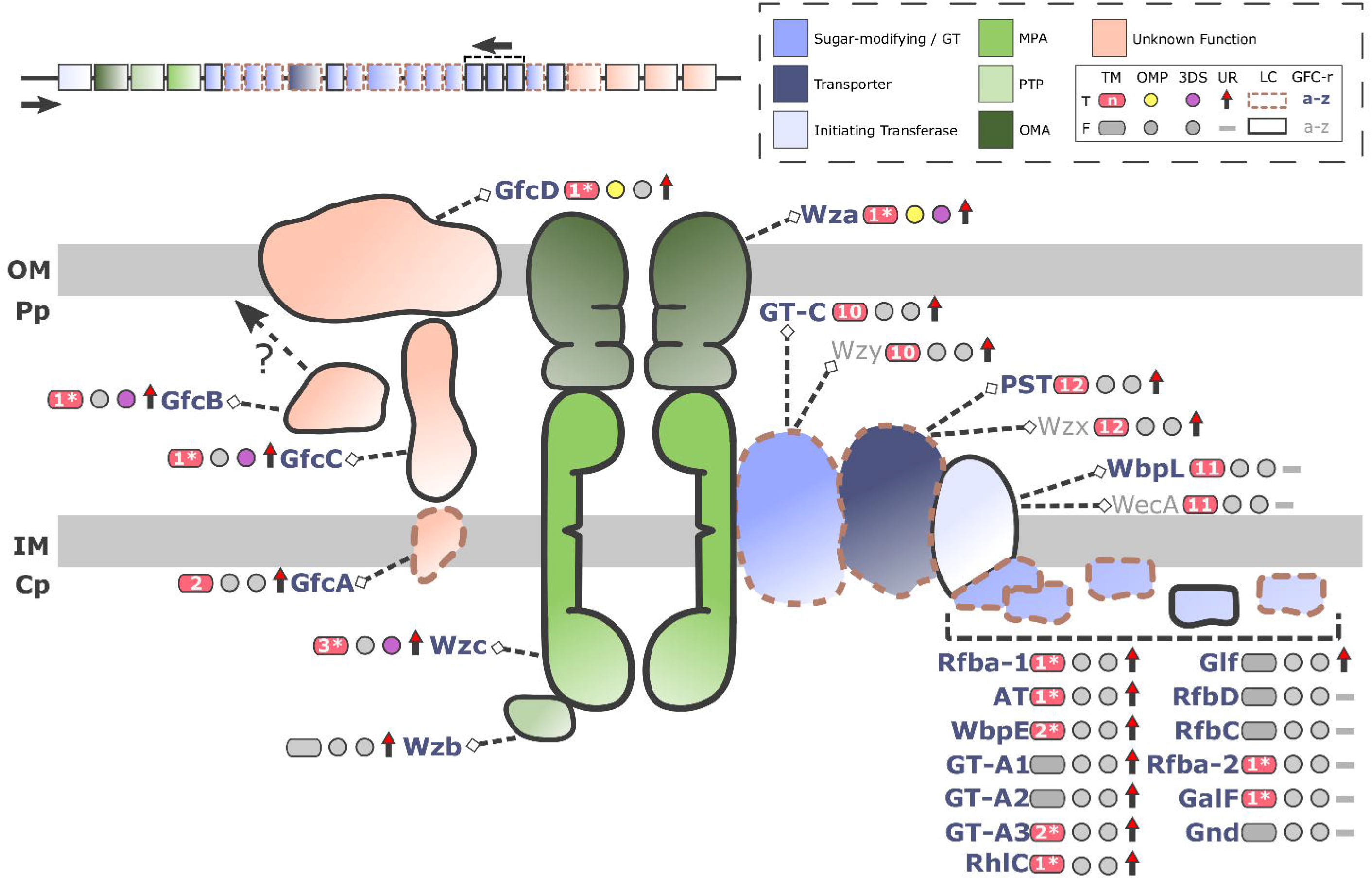
Schematic proposition of GFC machinery organization in *Pcb* 1692: The schematic view is based on the current model of Wza/Wzc polysaccharide exportation in group 4 capsule biosynthesis (Whitfield, 2006). Cell structure representations are abbreviated as follows: <u>o</u>uter-<u>m</u>embrane (OM), <u>p</u>eriplasm (Pp), <u>i</u>nner-<u>m</u>embrane (IM), and <u>c</u>yto<u>p</u>lasm (Cp). On top-right, the key-box show color assignments of known/unknown protein functionalities using the following abbreviations: <u>m</u>embrane-<u>p</u>eriplasmic <u>a</u>uxiliary protein (MPA), <u>p</u>rotein <u>t</u>yrosine <u>p</u>hosphatase (PTP), <u>o</u>uter-<u>m</u>embrane auxiliary protein (OMA). A binary table shows graphical representations for presence/true (‘T’) or absence/false (‘F’) of six parameters analyzed in the sequences following these abbreviations: transmembrane segments (TM), <u>o</u>uter-<u>m</u>embrane <u>p</u>rotein (OMP), available <u>3D</u> <u>s</u>tructure (3DS), significant infection-induced <u>u</u>p-<u>r</u>egulation in *Pcb* 1692 (UR), <u>l</u>ow <u>c</u>onservation amongst SRE (detailed below) assessed by synteny and sequence similarity (LC), genomic localization within the GFC region of *Pcb* 1692 (GFC-r). The number of detected TMs is represented within the pink shape: with an asterisk if the prediction methods diverge, or without asterisk if these methods converge in the number of TMs predicted. Low conservation (LC) is attributed to sequences exhibiting respectively less than 40 and 70 positive protein sequence and syntenic hits compared to 100 SRE (see Table S7 for details). On top-left, the genomic organization of GFC region is depicted using the features described in the top-right box. The arrows indicate the transcriptional orientation of the genes within the region.

By inspecting <u>o</u>uter-<u>m</u>embrane (OM) propensity (see ‘Experimental Procedures’) in sequences from the GFC region, only GfcD and Wza returned positive results (Supporting Information 6). This opens a question on the GfcB localization, which is generally regarded as an OM protein, but surprisingly could not be predicted as one (Fig. 5) (Zhou *et al*., 2006). The only structurally resolved encoded product from the *gfc* operon currently published is the GfcC, which is hypothesized to be a periplasmic protein (Sathiyamoorthy *et al*., 2011). Together the results presented here show that *Pcb* 1692 conserves canonical elements of Wza/Wzc system (i.e. *wzx/wzy*, and *wecA*) separated from the GFC region, which could support their encoded products as putative participants in the machinery. Additionally, evidence also indicates that PST, GT-4, and WbpL proteins encoded within the serotype-specific block of GFC region remarkably mimics structural features respectively found in Wzx, Wzy, and WecA, suggesting they could also undertake similar roles in the *wza/wzc* system (Fig. 5). Combined, these analyses expand the current view over GFC region gene-organization by applying extensive contextual comparison amongst SRE genomes, domain analysis and membrane topology predictions. The presence of serotype-specific blocks flanked by *wza-b-c* and *gfc* genes in SRE sheds light on a novel genomic architecture that combines the known pattern from group 1 capsule regions with typical genes involved group 4 capsule biosynthesis.

## 3. Concluding Remarks

In this article we presented an integrative approach that takes advantage of extensive public information to produce a reliable background for genome-wide studies in SREs organisms, including 100 strains from *Pectobacterium* and *Dickeya* genera. By combining this platform with an original transcriptome dataset, we shed light on strong genomic associations amongst virulence and interbacterial competition determinants in the SRE group, supported by coordinated transcriptional up-regulation of these elements in *Pcb* 1692 throughout disease development in potato tubers. Collectively, these findings provide strong evidence for gene associations and/or infection-induced transcriptional demand within important pathogenicity themes including PCWDE, T6SS, Ctv and the capsule biosynthesis machinery, which may pave the way for biotechnological applications in the near future.

## 4. Experimental Procedures

### Culture media, growth conditions and total RNA extraction

Wild type *Pcb* 1692 strain were grown on nutrient plate at 37°C for 16 hours. Overnight cultures were prepared by inoculating a single colony into 10 ml Luria-Bertani (LB) broth at 37 °C for 16 hours with constant shaking at 200 rpm. To obtain potato inoculated *Pcb* 1692 cells, healthy potato tubers (*Solanum tuberosum* cv. Mondial) were inoculated with *Pcb* 1692 (OD600 = 1) wild type strain as previously described in (Moleleki *et al*., 2017).The experiments were performed using three biological replicates, with three tubers per replicate. Macerated potato tissue was scooped out at 24 and 72 hpi and homogenized in double distilled water. Bacterial cells were recovered by grinding the scooped macerated potato tissues in 20 ml of double distilled water using autoclaved pestle and mortar. Starch material was removed by centrifuging ground tissue at 10000 rpm for 1 minute. Potato-inoculated and *in vitro* cultured bacterial cells were stabilized using the RNAprotect^®^ reagent (Qiagen, USA) according to manufacturer’s instructions. Total RNA from *in vitro* grown and potato-inoculated bacteria was extracted as previously described using the RNeasy mini kit (Qiagen, Hilden, Germany).

### Total RNA quality and cDNA library construction

The concentration and purity of each extracted total RNA sample was evaluated using spectrophotometric analysis (NanoDrop^®^ ND-1000; NanoDrop^®^ technologies, Wilmington, DE) at a ratio of 230/260 nm. Using the Agilent 2100 Bioanalyzer (Agilent Technologies, Inc.), total RNA samples’ concentration, RIN and 28S/18S ratio were determined. 200 ng aliquots of total RNA from *Pcb 1692* extracted from *in vitro* grown cells (16 h) and infected potato tubers at 24 and 72 hpi was used to prepare cDNA libraries. The Illumina sequencing service was provided by the BGI Co., Ltd (China). TruSeq RNA Sample Prep Kit v2 (Illumina, USA) was used to construct the cDNA libraries following the manufacturer’s protocol. In summary, the mRNA was cleaved into small fragments, followed by the synthesis of first-strand cDNA with random hexamer-primed reverse transcription. RNase H and DNA polymerase I were used to synthesize second-strand cDNA. The double-stranded cDNA was subjected to terminal modification, by addition of adenosine and ligated with adapters. Adaptor-ligated fragments with suitable sizes were selected and enriched by PCR using the PureLink_TM_ PCR Purification Kit (Invitrogen, USA) to create the cDNA libraries for sequencing. The paired-end sequencing (PE91) was performed on the Illumina HiSeq 2000 sequencing platform. The data have been deposited in NCBI’s Gene Expression Omnibus (GEO) and are accessible through the GEO accession number, GSE102557.

### Reads mapping, gene expression and statistical analysis

Quality assessment of the raw data was performed by fastqc software (https://www.bioinformatics.babraham.ac.uk/projects/fastqc), then low quality regions were trimmed by Trimmomatic v 0.36 (Bolger *et al*., 2014). The trimmed RNA-Seq reads were mapped to the *Pcb* 1692 reference genome (GCF_000173135.1) using hisat2 v 2.1.0 (Kim *et al*., 2015). Raw read-counts were performed in R environment (https://www.r-project.org/) by the *featureCounts* package (Liao *et al*., 2014) and subsequent statistical analysis of differential expression by edgeR package (Robinson *et al*., 2010). The threshold parameters used to assign differential expression were FDR < 0.01 (Benjamini & Yekutieli, 2005), and absolute log2fold-change > 1 (up-regulated), or log2fold-change < −1 (down-regulated). Genes’ transcriptional profiles were graphically rendered by Gitools (Perez-Llamas & Lopez-Bigas, 2011).

### Orthology analysis, domain architectures detection, and gene neighborhood screenings

Orthology relationship between protein sequences was assessed using the OrthoMCL pipeline (Li *et al*., 2003), which takes tabular results from a Blastp search under specific parameters (options: -seg yes -outfmt 6 -num_threads 3 -num_alignments 100000 -evalue 1e-05) as input. The sequences are clustered in orthologous groups labeled with “OG” numeric tags (e.g. OG_1; OG_2). OrthoMCL was performed with MCL granularity of 1.5. In parallel, all sequences were characterized by using HMMER3 (Eddy, 2011) supported by Pfam-A database (Finn *et al*., 2010) in order to generate in-house predictions of conserved domain architectures through Hidden Markov Models (HMM) profiles (Baum & Petrie, 1966). Orthology and domain architecture information are then combined with genomic coordinates from each genome by custom Perl scripts in order to generate annotated gene neighborhood screenings.

### Ectopic expression of predicted T6SS-dependent toxins

Removal of Rat FABP1 gene from pTrc99A (AddGene) removed some cut sites from the MCS. *KnpI* and BamHI cut sites were reintroduced into the plasmid by incorporation of the cut sites into the 5’ region of the PCR primers (H5907_F/R) (Table S8). PCR amplification of *PCBA_RS05790* (hypothetical protein upstream of WHH-containing nuclease) incorporated *XbaI, KpnI*, and BamHI upstream of the ORF and BamHI and *SalI* downstream of the ORF. The PCR product was subcloned into pJET1.2/blunt and excised with *XbaI/SalI* restriction digest. pTrc99A (without rat FABP1) was digested with *XbaI/SalI* and ligated with the excised PCR product. The gene was removed using BamHI digest and the linear plasmid re-ligated. In this manner BamHI and KnpI were reintroduced into the MCS to facilitate cloning of effector genes. The resulting plasmid is referred to as pTrc100 as its MCS differs slightly from pTrc99A, but the rest of the plasmid remains unchanged.

Effector genes (*PCBA_RS05785:* WHH-containing nuclease, SacI/PstI; *PCBA_RS05775:* D123-containing protein, *KpnI/PstI* digest; *PCBA_RS18045:* phospholipase, SacI/PstI digest; and *PCBA_RS22965:* AHH-containing nuclease, SacI/PstI digest) were ligated into arabinose-inducible expression plasmid pCH450; immunity genes (*PCBA_RS22965*; AHHi; KpnI/PstI digest) were ligated into IPTG-inducible pTrc100, and respectively transformed into *E. coli* DH5-alpha. Overnight cultures were grown in 0.1% glucose to repress expression from the arabinose-inducible promoter. Cultures were washed in 10 mM MgSO4, adjusted to an OD600 of 0.05 in fresh LB broth supplemented with 5 μg/ml tetracycline, 50 μg/ml ampicillin, and 1 mM IPTG, as required. After 30 minutes, expression from pCH450 was induced with 0.2% L-arabinose. Where necessary, expression from pTrc100 with IPTG was induced immediately. Optical density (600 nm) was measured hourly.

### Collinearity analysis, prophage-origin predictions and capsule regions inspection

In order to predict collinear conservation in SRE genomes, we first obtained genomic annotation and protein sequence data from all *Pectobacterium* and *Dickeya* strains available on RefSeq database (https://www.ncbi.nlm.nih.gov/refseq) (Table S9) until December/2017, totaling 100 strains including *Pcb 1692*. We then performed the synteny analysis using MCScanX (Wang *et al*., 2012) allowing minimum match size of two genes to call syntenic blocks (option: -s 2). MCScanX takes the tabular results of a non-stringent Blastp search (Altschul *et al*., 1990) as input (option: -evalue 1). Total number of positive matches of gene-products (from one organism) against all other SRE protein datasets resulting from both Blastp and MCScanX were parsed by in-house Perl scripts (https://www.perl.org/). Graphical ideograms were scripted in Circos (Krzywinski *et al*., 2009) compiling syntenic regions in the genomes as ribbon-links between chromosomes, as well as Blastp and MCScanX hit counts from pairwise comparisons previously described. Prophage-origin predictions were conducted by using PhiSpy software (Akhter *et al*., 2012), in parallel with Blastp search support (options: -qcov_hsp_perc 40 -evalue 1e-05) against a Phantome (http://www.phantome.org), Phast (Zhou *et al*., 2011) and NCBI bacteriophage-sequence databases. In parallel we have performed genome-wide synteny predictions on *Pcb* 1692 and *Pcc* strain BCS2 using recently predicted prophage regions published by Varani *et al*. (2013) . The capsule regions were initially surveyed using HMM-profile search (Eddy, 1998) based on the curated database of capsule-related domains made public by Rendueles *et al*. (2017) . Next, the *Pcb* 1692 genome was programmatically scanned for segments harboring at least three consecutive matches carrying the above-mentioned capsule-related domains with two non-matches gap allowed for greater sensitivity. These segments were then manually inspected in an effort to identify detectable functionality in the adjacent genes that could support the existence of a capsule region.

### Sequence search, alignment and membrane topology predictions of GfcA-related entries

The PCBA_RS09180-related sequences were obtained using HHblits online tool by default parameters (Remmert *et al*., 2011). HHblits positive hits were filtered by at least 40 matched amino-acid columns in HMM-HMM alignment, representing ~40% of the query (PCBA_RS09180) sequence (Table S7). Next all above-threshold entries were retrieved. Out of 50 above-threshold entries, 27 were supported by publicly available genome-wide data, making them suitable for gene neighborhood screening. Genomic data for these 27 structures (complete genome/scaffold/contig) were then obtained from RefSeq database for gene neighborhood screening. CDART database were inspected in order to obtain extensive representativeness of YjbE, YjbF, Caps_synth_GfcC, and YjbH (PFAM: PF11106, PF11102, PF06251, PF06082). (Geer *et al*., 2002). Sequence alignments were performed using Clustal Omega (Sievers & Higgins, 2018). Prediction of transmembrane segments were conducted by using TMpred (Hofmann, 1993) and TMHMM (Moller *et al*., 2001) online servers in parallel. Signal peptides were predicted by SignalP 4.1 server (Emanuelsson *et al*., 2007) and Signal-Blast (Frank & Sippl, 2008). Detection of putative outer-membrane proteins was conducted by using HHomp (Remmert *et al*., 2009).

### STRING network integration

In order to integrate *Pcb* 1692 with STRING database (von Mering *et al*., 2005), orthology annotations with *Escherichia coli* strain K-12 and *Pseudomonas aeruginosa* strain PAO1 were used. The *Pcb* 1692 entries conserving orthology with the model organisms were supplied to STRING correlational database. This provided association between annotated orthologs with the respective Gene Ontology (GO) terms (Carbon *et al*., 2009) for each sequence.

## 5. Acknowledgements

This research study was funded by the National Research Foundation (NRF), South Africa through Competitive Funding for Rated Researchers (CFRR 98993); NRF Bioinformatics and Functional Genomics (BFG 93685) and NRF Research Technology and Transfer Fund (RTF) 98654. DYS NRF BFG Post-Doctoral Fellowship, DB-R University of Pretoria Post-Doctoral Fellowship. CKT PhD Bursary was funded by the University of Pretoria Bursary. Any opinion, findings, conclusions or recommendations expressed in this material is that of the author(s) and the NRF does not accept any liability in this regard.

## 6. Author Contributions

Conception or design of the study: D.B.-R., C.K.T.; and L.N.M.; Acquisition, analysis, or interpretation of the data: D.B.-R., C.K.T.; N.M., D.Y.S., S.K., and L.N.M.; Wrote the manuscript: D.B.-R., N.M., and L.N.M.

## 7. Abbreviated Summary

Through large-scale gene expression obtained from *Pcb* 1692 and comparative genomics using 100 *Pectobacterium* and *Dickeya spp* this report uncovers association amongst key pathogenicity themes in the main Soft-Rot *Enterobacteriaceae* taxa. The approach enabled identification of striking transcriptional and contextual genomic association between: (i) the WHH/SMI1_KNR4 toxin/immunity pair with type VI secretion system, (ii) the carotovoricin prophage with type I secretion system, and (iii) a serotype-specific block ranging 13-27 genes with characteristic group-4-capsule genes.

